# Niche Overlap Is Not Enough: Same Overlap, Opposite Dynamics

**DOI:** 10.1101/2025.09.05.674242

**Authors:** Akiva Goldberg, Oshrit Shtossel, Yoram Louzoun, Nadav M. Shnerb

**Affiliations:** Department of Physics, Bar-Ilan University, Ramat Gan 52900, Israel; Department of Mathematics, Bar-Ilan University, Ramat Gan 52900, Israel

## Abstract

Niche overlap (NO) is a central concept in coexistence theory, determining the outcome of competitive interactions at equilibrium. Yet, NO is a composite quantity that collapses multiple mechanisms and thus cannot by itself capture community dynamics. We quantify this gap by analyzing one of the simplest out-of-equilibrium observables: temporal correlations in species abundances, widely used in microbial ecology. Within a generalized consumer-resource framework, we separate the mechanistic traits that shape dynamical responses and show that systems with identical NO can display opposite correlation patterns. The decisive predictor is the yield–depletion mismatch (YDM) - the mismatch between species’ depletion and yield dissimilarities. Analytical theory, stochastic simulations, and a reanalysis of microbial time-series consistently show that YDM, not NO, sets the sign and magnitude of abundance correlations, whereas instantaneous growth-rate correlations increase with NO. Our results suggest that other out-of-equilibrium properties, such as invasion probabilities and coexistence robustness, will likewise be largely independent of NO and instead controlled by distinct out-of-niche parameters.

## I. INTRODUCTION

The concept of niche overlap, or conversely, niche segregation, lies at the heart of ecological coexistence theory. As niche overlap between populations increases, so does the extent to which they limit each other’s abundance, making coexistence increasingly dependent on compensating fitness differences [1, 2]. Consequently, considerable effort has been devoted to defining niche overlap within theoretical models and quantifying it in empirical and experimental systems [3, 4]. Beyond its central role in coexistence theory, quantifying niche overlap also enables the partitioning of a diverse community into functional groups, which can facilitate the interpretation and analysis of community dynamics [5, 6].

Despite its key role, it remains unclear to what extent niche overlap (NO) alone can account for the dynamical properties of ecological communities, even in the absence of fitness differences. NO is a composite parameter that summarizes the net competitive effect among species at equilibrium, but different underlying mechanisms can lead to the same NO value. Thus, two communities with identical NO might, in principle, respond very differently to perturbations. This raises the question of whether NO is sufficient to characterize out-of-equilibrium behavior, or whether additional mechanistic parameters are required [7, 8].

One of the most accessible and analytically tractable manifestations of out-of-equilibrium dynamics is the pattern of correlations between species’ abundances over time. Such correlations quantify the extent to which fluctuations in one species’ abundance are mirrored by fluctuations in another’s, and they can be estimated from time-series data. Recently, abundance correlations have received considerable attention, particularly in microbial community studies, where high-resolution time-series are increasingly available [5, 9, 10]. These correlations are often interpreted as indicators of niche overlap (NO), under the assumption that species with greater overlap will display stronger correlations. However, even at the *a priori* level, abundance correlations expose a limitation of NO as a sole descriptor. Even if correlations increase with NO [5], the direction of the effect is intrinsically ambiguous: higher overlap aligns species’ responses to environmental fluctuations (favoring positive correlations) while simultaneously intensifying competitive coupling (favoring negative correlations). Consequently, the sign and magnitude of abundance correlations cannot be inferred from NO alone, motivating the out-of-niche metric developed below.

To address this conundrum, we consider a consumer–resource (CR) system [11], where species interact indirectly through shared resources. In such systems, each species is characterized by two distinct functional profiles. Its *yield* quantifies the growth benefit it gains from one unit of resource, i.e., how efficiently resource uptake translates into population growth. Its *depletion* measures the rate at which it consumes or removes each resource per unit of its own biomass. At equilibrium, or when resource dynamics are much faster than the consumer dynamics, these two profiles combine to produce the effective competition coefficients between consumers, and hence the niche overlap (NO). Crucially, the mapping from the full set of CR parameters to the reduced consumer–consumer competition matrix is highly redundant: many distinct CR configurations yield the same NO and the same equilibrium state, yet they may differ in their out-of-equilibrium behavior. This redundancy enables keeping NO fixed while systematically varying the underlying yield and depletion traits.

The conflicting expectations that arise from NO motivate the search for an alternative predictor. We show that the decisive predictor for the correlations is the *yield–depletion mismatch* (YDM) between species, i.e., the difference between their depletion dissimilarity and their yield dissimilarity. Analytical calculations, stochastic simulations, and reanalysis of microbial time-series data [5] consistently reveal that YDM determines both the sign and magnitude of abundance correlations. Niche overlap itself plays no direct role in these patterns.

The finding that abundance correlations are governed by yield–depletion mismatch (YDM) rather than niche overlap (NO) underscores a broader principle: out-of-equilibrium properties of ecological communities can hinge on mechanistic parameters that are invisible to the reduced competition matrix used to define NO. This perspective suggests the need for a more comprehensive framework in which niche overlap reflects only the static equilibrium structure, while additional “out-of-niche” parameters capture the dynamical responses of communities. We return to this point in the Discussion, where we outline the implications of this broader view for theories of community assembly and stability.

## II. METHODS

### A. Model Framework

Our starting point is a slight modification of MacArthur’s consumer–resource model for *S* consumers (*n*_*i*_(*t*) is the instantaneous abundance of the *i*-th consumer species) and *Q* resources (where *R*_*k*_(*t*) is the available biomass of the *k*-th resource),

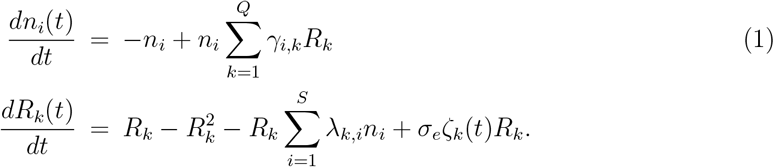

Here *γ*_*i,k*_ is the *{i, k}* element of the *S×Q* matrix Γ that quantifies the yield, i.e., the increase in growth rate of the *i*-th consumer due to the consumption of the *k*-th resource, whereas *λ*_*k,i*_, the elements of a *Q × S* matrix Λ, quantify the depletion of the resource due to the consumer. The *σ*_*e*_ terms represent stochastic variations in the growth rate of the resources, where *ζ*_*k*_(*t*) is a white-noise process.

Our aim is to obtain a reduced model in which niche overlap can be defined in the simplest and most transparent way. To this end, we consider the deterministic case (*σ*_*e*_ = 0) and assume a quasi-steady state (QSS), which allows us to integrate out the resource dynamics by setting *dR*_*k*_*/dt* = 0 (see Supplementary A). With a suitable choice of parameters, this procedure yields a Generalized Lotka–Volterra equation for the consumers of the form

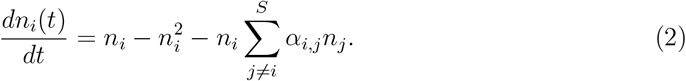

In this reduced form, all species share the same intrinsic growth rate (set to one), and in the absence of competition (*α*_*i,j*_ = 0 for all *i ≠ j*) they also share the same carrying capacity (set to one). These simplifications do not affect the generality of the principles demonstrated below; instead, they guarantee that the coefficients *α*_*i,j*_ directly quantify the niche overlap between species *i* and *j* [3].

Eqs. (1) are mapped, when *σ*_*e*_ = 0, to Eqs. (2), provided that each row of the Γ matrix sums up to 2, and that the interaction matrix satisfies ***α*** = ΓΛ, with *α*_*i,i*_ = 1 (see Figure 1). The elements of ***α*** quantify the classical niche overlap (NO) between species [3] and uniquely determine the consumer equilibrium density (see Supplementary A). In contrast, as we shall see, its relevance to the magnitude of the correlations is at most secondary and indirect.

**FIG. 1:**
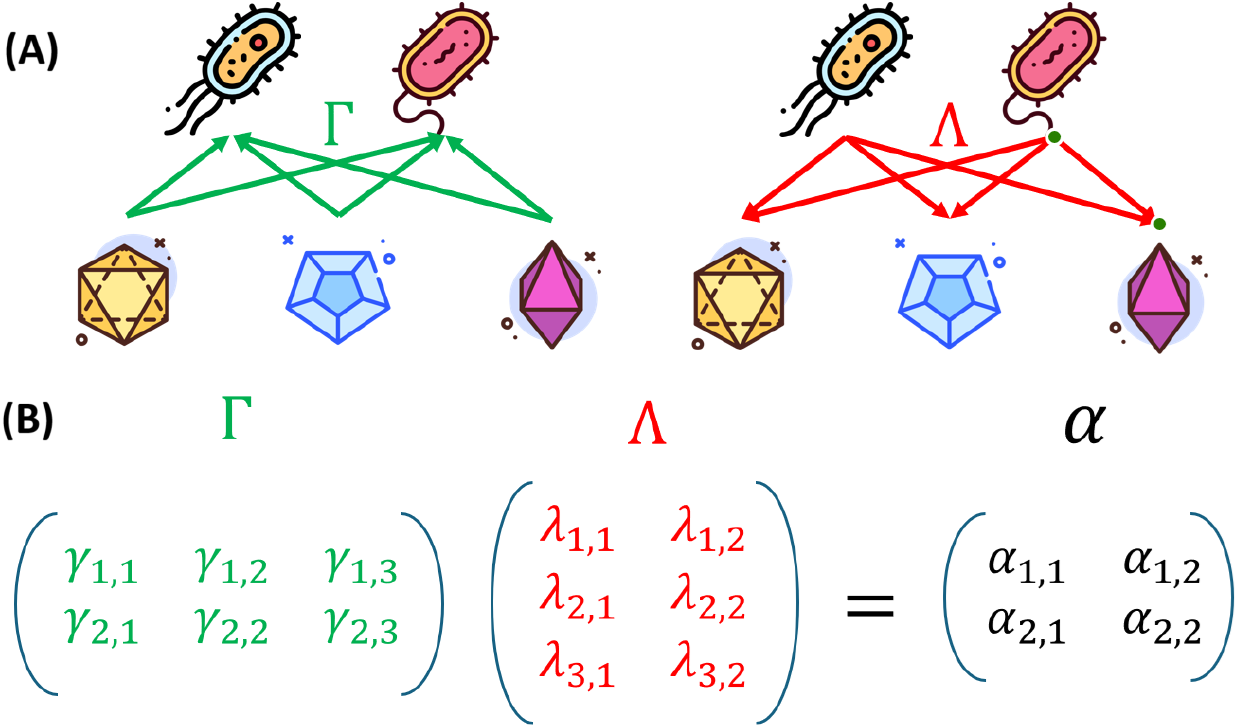
The consumer-resource model, Eq. (1), describes the dynamics of *S* species and *Q* resources. Panel (A) illustrates the case in which *S* = 2 and *Q* = 3. Each elements of the Γ matrix, *γ*_*i,k*_, describes the yield (growth rate per unit mass) of the *i*-th species due to the *k* resource (green arrows). Element of the Λ matrix, *λ*_*k,i*_, quantifies the depletion of the *k* resource due to the presence of the *i*-th consumer species. Panel (B) illustrates the mapping of the consumer-resource model (1) into a Lotka-Volterra model for the consumer species when the nutrient dynamic is fast, so one imposes *dR*_*k*_*/dt* = 0. To get Eq. (2) in that specific parametrization each row in the Γ matrix must sum up to 2, and then the ***α*** matrix is obtained as the product of Γ and Λ. It is important to emphasize that this mapping is highly redundant; that is, there are many possible choices of the Γ and the Λ matrices that correspond to the same *α*.

In the parameterization we used, the consumer–resource model has 2*SQ* parameters (the two matrices *γ* and *λ*), which are constrained by *S*^2^ + 2*S* conditions (the matrix ***α*** and the row sum of Γ). Therefore, as explained in detail in Supplementary **??**, the mapping is highly redundant, with many CR systems give rise to the same Lotka–Volterra system. We exploit this redundancy to investigate how consumer abundance correlations vary across consumer-resource systems that share the same ***α***, and to examine how such correlations depend on the underlying mechanistic processes that are not captured by the effective interaction coefficients alone. The core numerical task is to generate the two matrices randomly, up to the imposed constraints. In Supplementary **??**, we detail the two sampling strategies employed in this work.

### B. Calculating correlations

We employed three complementary methods to calculate abundance and growth-rate correlations between pairs of species. First, we directly integrated the stochastic differential equations (1) numerically, tracking abundance and growth rates over time. Second, we identified the deterministic fixed point and estimated the expected correlations using linear response theory, based on the Jacobian matrix at equilibrium and the noise covariance structure. This method is semi-analytical, as it involves inverting the Jacobian matrix. Finally, in specific cases involving two species and two resources with carefully chosen interaction coefficients, we were able to solve the system analytically and obtain exact expressions for the correlations. These analytical results guided us in formulating an informed conjecture for a general metric that predicts the sign and magnitude of abundance correlations. The technical details of all three methods are provided in Supplementary B and C.

### C. Empirical Data Analysis

We reanalyzed the publicly available dataset from Crocker *et al*. [5], who monitored the abundance dynamics of a 20-strain synthetic microbial community over nine batch-growth cycles in 32 distinct environments, each defined by a unique mixture of carbon sources. Raw abundance time-series and monoculture measurements of growth rate and resource depletion were obtained from the dataset deposited in their public repository.

Based on these data, we computed the functional dissimilarities in yield and depletion and evaluated the correlation between abundance time series for each pair of strains across environments. Specifically, we calculated Spearman’s rank correlations over the history of each environment, and averaged the resulting values across all environments in which both strains were sufficiently abundant. These empirical correlations were then compared to theoretical predictions based on the YDM *D*.

## III. RESULTS

### A. The yield-depletion mismatch parameter *D*

We would like to analyze the dynamics of the consumers when stochasticity is introduced into the time evolution of the resources. Our primary goal is to assess the relationships between NO and an out-of-equilibrium feature, the temporal abundance correlations between coexisting species. We will show that NO does not determine correlations, and we will develop an alternative metric that is useful for practical applications and aids intuition.

Specifically, the behavior of the consumer subsystem is studied when the resources experience environmental stochasticity, i.e., when *σ*_*e*_ *>* 0 in Eq. (1). Stochastic fluctuations affecting each resource are assumed to be mutually independent, ⟨*ζ*_*k*_(*t*), *ζ*_*ℓ*_(*t*)⟩ = 0 if *k≠ ℓ*. Nevertheless, when two consumer species depend on the same resource, fluctuations in that resource are expected to induce correlated responses in their abundances. The aim here is to identify the parameter controlling these correlations in the stochastic regime.

If two consumer species, *i* and *j*, have identical yield and depletion profiles, that is, if for all *k, γ*_*i,k*_ = *γ*_*j,k*_ and *λ*_*k,i*_ = *λ*_*k,j*_, then they are dynamically indistinguishable, even if they differ biologically in superficial ways (e.g., coloration, synonymous mutations). The relevant functional differences between the two species are quantified by the differences between their *γ*s and their *λ*s. We sought to examine whether, and how, the correlations in species abundances depend on these functional differences. To operationalize these functional differences, we introduce the *Yield–Depletion Mismatch* (YDM) metric *D*, based on yield and depletion matrices, through,

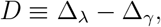

where Δ_*γ*_ = 1 − cos *θ*_*ij*_(*γ*) is the cosine distance between the *row* vectors 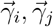 of the yield matrix Γ, and Δ_*λ*_ = 1−cos *θ*_*ij*_(*λ*) is computed from the *column* vectors 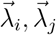 of the depletion matrix Λ. Thus, *D* depends directly on Γ, Λ (and not only on *α* = ΓΛ).

The metric *D* is a *theory-guided conjecture* grounded in an exact solution of a solvable two-species/two-resource case that yields closed expressions for correlations. In that case, as detailed in Supplementary C, *D* captures the shape, zero-crossings, and the peak of the correlation. Moreover, *D* is a bounded, normalized parameter: since 0 ≤ Δ_*γ*_, Δ_*λ*_ ≤ 2, it follows that −2 ≤ *D* ≤ 2; consequently, *D* is scale-free and enables consistent comparisons across systems.

These numerical experiments reveal that *D* is, indeed, strikingly predictive, consistently recovers the salient features of abundance correlations without additional fitting. Figure 2 illustrates this agreement: predictions based on *D* closely tracks the measured correlations across parameter sweeps and model variants, underscoring *D*’s practical value as a compact, mechanistic predictor.

**FIG. 2:**
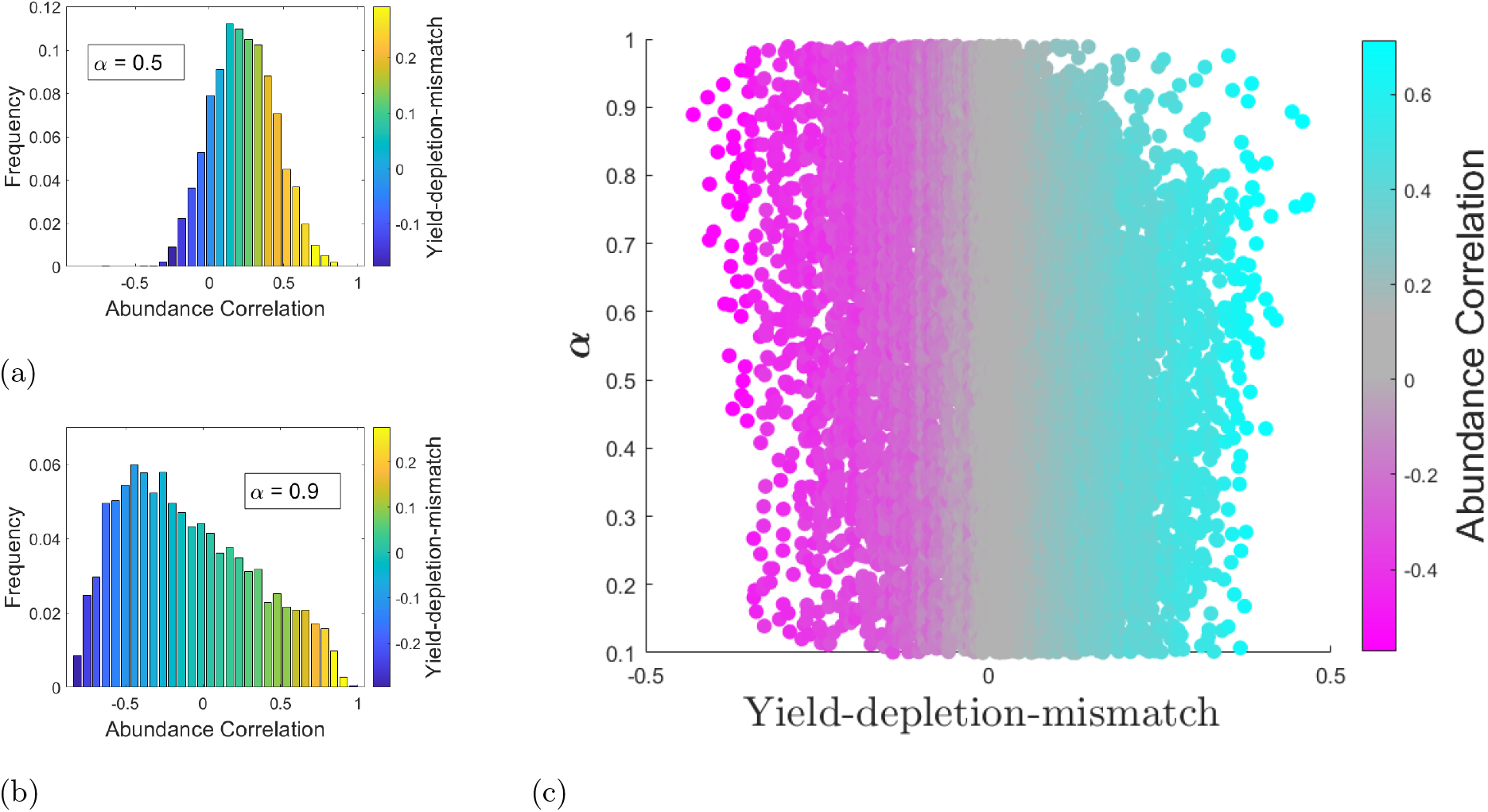
Inadequacy of niche overlap *α*, and effectiveness of the yield-depletion mismatch (YDM) *D*, as predictors of abundance correlations. The histograms in panels (a) and (b) show the frequency of abundance correlations between two consumer species for many realizations with *α* = 0.5 (a) and *α* = 0.9 (b). The color of each bar indicates the mean value of *D* in that bin. We observe that *α* alone is a poor predictor of the correlation, whereas the YDM *D* captures it well. Panel (c) illustrates this result in a more general setting: here, color indicates the level of correlation obtained from numerical experiments across the *D*–*α* plane. Clearly, correlations are a function of *D*, not of *α*, although the relative frequency of negative *D* values in the sample space increases as *α* → 1. We first fixed a specific *α* in a two-consumer, three-resource model, and then used the redundancy of the consumer–resource model to generate many randomly chosen Γ and Λ matrices that yield the same *α*, as described in Supplementary A. For each realization, we computed the correlation between the abundances of the two consumer species, with stochasticity entering through the resource dynamics as described in Eq. (1).

The basic ecological insight captured by the yield–depletion mismatch *D* is illustrated in Fig. 3. The two limiting cases, Δ*γ* ≫ Δ*λ* and Δ*γ* ≪ Δ*λ*, correspond respectively to strongly negative and strongly positive abundance correlations, as explained in the caption.

**FIG. 3:**
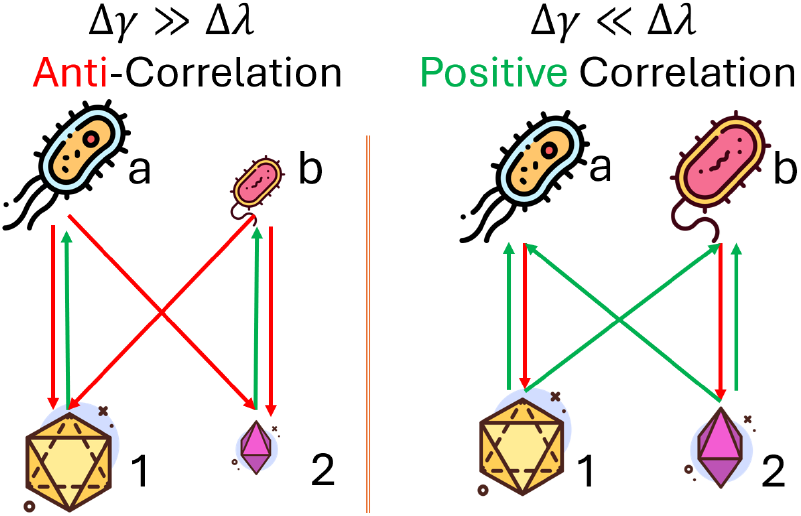
Intuition Behind the Yield–Depletion Mismatch *D*. Two schematic examples corresponding to limiting cases. Green lines represent yield; red lines represent depletion. **Left:** Δ*γ* ≫ Δ*λ*, i.e., each consumer depends on a different resource (green arrows) but depletes (red arrows) both in a similar way. An increase in one resource strongly benefits only one species; since depletion is not species-specific, this harms the other species, producing negative correlations. **Right:** Δ*γ* ≪ Δ*λ*. Now, both consumers are generalists and respond similarly to increases in any resource. Although their depletion profiles differ, the shared response to environmental variation dominates, producing positive correlations.

In Fig. 4, we show that *D* is applicable as a predictor of correlations even when the number of resources is increased, or when the community includes many consumer species. Note that we considered only pairs of species for which the deterministic dynamics allow for a stable coexistence. Other cases, such as those in which the system transitions to chaotic dynamics [12, 13], or cases involving transient species that persist solely due to immigration, require a separate discussion.

**FIG. 4:**
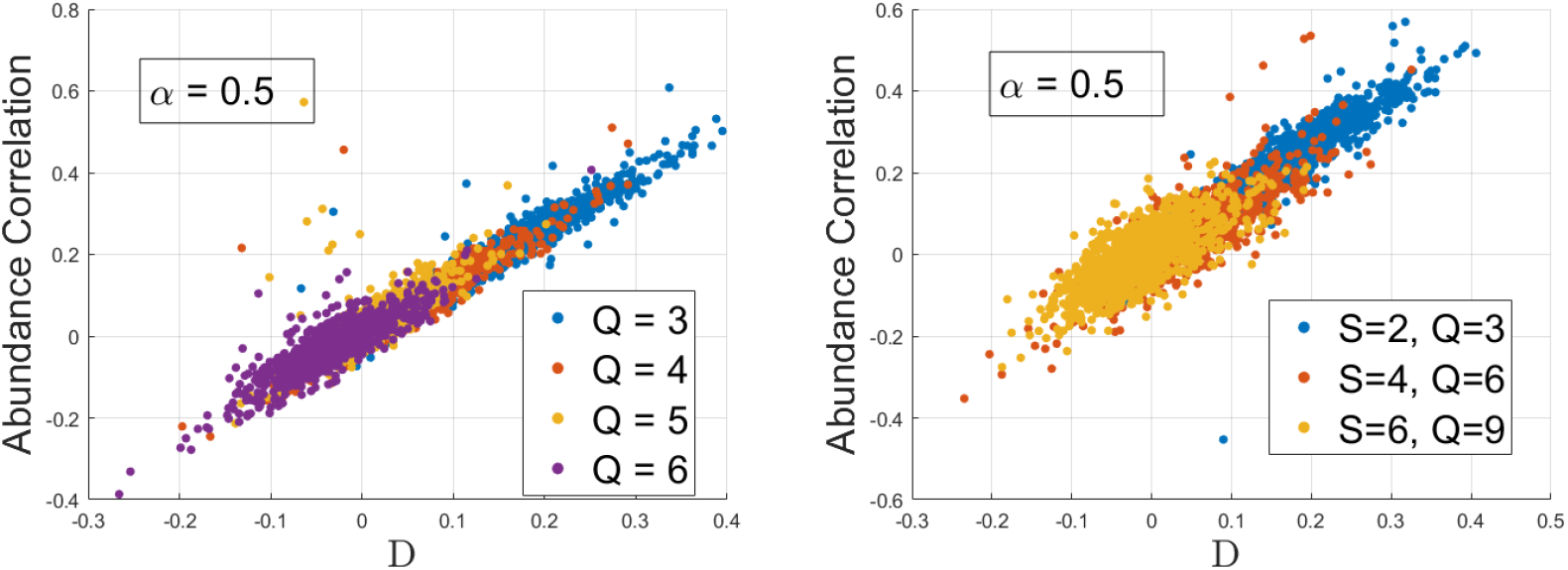
Effect of more resources and more species. The left panel illustrates the correlations vs. *D* for a fixed value of the niche overlap *α* = 0.5, in a two-species model with a varying number of resources. Clearly, the credibility of the YDM parameter *D* as a predictor of correlations is not affected by the number of resources. The right panel shows the same outcomes for communities with different numbers of species and different numbers of resources.

### B. The time-averaged neutral limit

The results shown in Fig. 2 suggest that the YDM parameter *D* uniquely determines the abundance correlation between consumer species. However, we also observe that when the system is, on average, almost neutral, i.e., as *α* → 1, it becomes increasingly difficult to find Γ and Λ matrices for which *D* takes positive values. In such cases, the resulting correlations are typically very close to minus one, indicating near-perfect anticorrelation. This result is relevant to many recent works on time-averaged neutral dynamics [14–17].

One could hypothetically attribute this pattern to imperfect sampling of the underlying parameter space. When working at fixed *α*, some entries of the Γ and Λ matrices are constrained, while the remaining parameters are randomly sampled. It is difficult to guarantee uniform coverage of the entire feasible matrix space under such constraints. Nevertheless, we believe the observed trend is genuine. In Fig. 5, we compare the average correlations resulting from two different sampling methods for the matrix space (detailed in Supplementary A). As far as can be judged from these results, the approach to the limit of neutral consumer dynamics (i.e., an ***α*** matrix whose all entries equal to one) appears to be associated with strongly anti-correlated abundance fluctuations of consumer species.

**FIG. 5:**
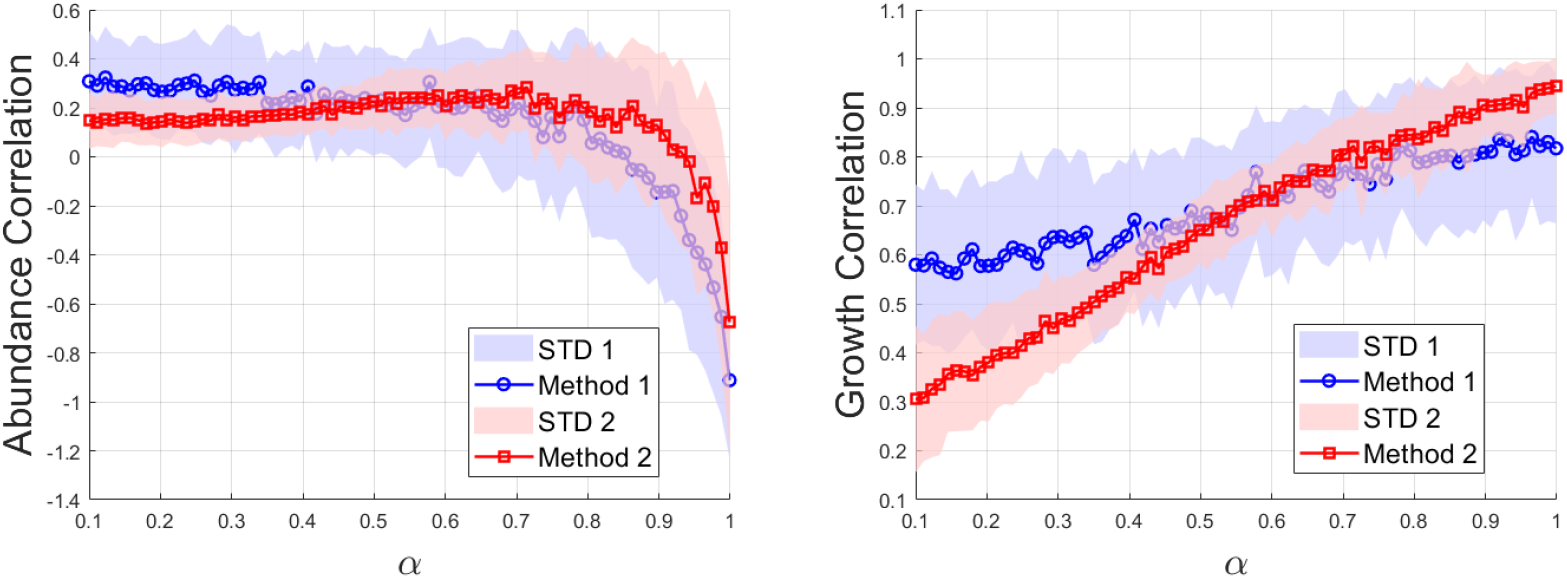
Correlations vs. niche overlap. the mean (markers) and the standard deviation (shaded area) of the correlation values are plotted against niche overlap *α* for two-species, three-resources dynamics. The correlations in abundance (left panel) are quite a bad indicator for niche overlap - they are almost independent of *α* until *α >* 0.8, say, and then drop towards (−1) at the “neutral” limit *α* → 1. On the other hand, the correlations in growth rate (right panel) are proportional to niche overlap. The space of Γ and Λ matrices was sampled by two different methods, as explained in supplementary A. While these two methods yield different quantitative results, the qualitative picture is more or less the same.

### C. Growth-rate correlations

Abundance correlations fail as indicators of niche overlap. By contrast, correlations in growth rates (Figure 5, left panel) provide a much better proxy: competitive effects are relatively small, and the shared response to environmental fluctuations dominates.

At first glance, this result may seem surprising. Since abundance is obtained by integrating the instantaneous growth rates, one might expect abundance correlations to track growth-rate correlations. Yet the two can diverge: growth rates may be positively correlated while the corresponding abundance time series are negatively correlated. Indeed, it is easy to construct functions with this property; for example,

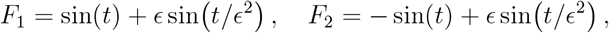

which satisfy it in the limit *ϵ* → 0.

### D. Empirical support

Empirical validation of these theoretical results requires abundance data for several competing species, obtained under conditions where the concentrations of relevant metabolites fluctuate while other environmental aspects (such as temperature and pH) are held constant. In addition, monoculture assays are needed to determine each species’ yield and depletion rate per metabolite.

While such comprehensive datasets are rare, a recent study provides a close approximation. Crocker *et al*. [5] tracked the temporal dynamics of a synthetic community composed of 20 microbial strains. Each of these strains was first grown in monoculture on one of 10 possible carbon sources (glucose, glycerol, etc.), and its growth rate and its depletion level were measured (see Supplementary D). Subsequently, the full community was introduced into an environment containing a random mixture of these 10 carbon sources and subjected to 9 batch growth cycles, with a 10-fold dilution into fresh media (with the same carbon composition) at the end of each cycle. A total of 32 such experiments were conducted, each with a different carbon-source composition, yielding a dataset of species abundances for 20 strains over 9 dilution cycles across 32 environments.

For each pair of species, and for each carbon-source composition (i.e., each environment), we thus have 9 simultaneous abundance measurements and can compute the correlation between the corresponding time series. Although metabolite concentrations are largely reset after each dilution, fluctuations do occur from batch to batch. Moreover, there are also changes in metabolite levels during each growth cycle due to consumption by other species, so it is reasonable to expect that at least part of the observed correlation reflects fluctuations in metabolite availability. Another part presumably reflects fluctuations in environmental factors such as temperature and pH, for which we do not have direct measurements.

To test whether our theoretical predictions hold in a real microbial system, we analyzed the dataset described above by computing, for each pair of species, the mean abundance correlation across all 32 environments and examined its dependence on two functional dissimilarity metrics: the yield-depletion mismatch *D* = Δ*λ* − Δ*γ*, and the total functional dissimilarity *D*_*T*_ = Δ*λ* + Δ*γ*. As shown in Fig. 6, a statistically significant positive trend is observed between the correlation and *D*, whereas the trend with *D*_*T*_ is weak and not significant. This suggests that the asymmetry between depletion and yield, rather than the total functional dissimilarity, is more relevant for predicting co-abundance patterns, in line with our theoretical framework.

**FIG. 6:**
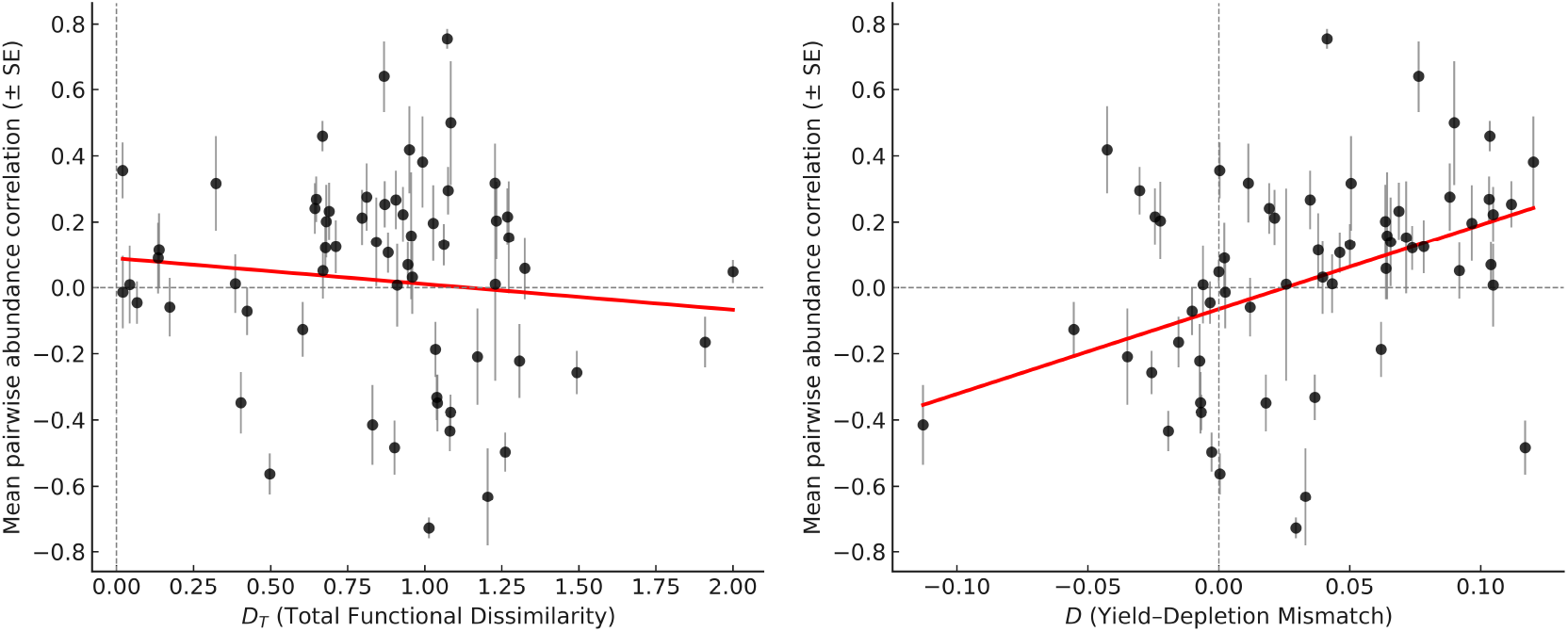
Abundance correlations in empirical community. Abundance correlations between microbial strains as a function of total functional dissimilarity *D*_*T*_ (left, Panel A) and the YDM parameter *D* (right, Panel B). To avoid the influence of inferior species approaching extinction, we included in the analysis only the ten most abundant species in each environment. To obtain meaningful confidence intervals, we further restricted the analysis to species pairs that co-occurred among the top ten in at least three environments. In addition, we considered only data from the second day onward to minimize the effect of transients. A statistical robustness checks of our results are presented in Supplementary D. In the left panel, the observed downward trend is not statistically significant (*p* = 0.46), whereas in the right panel, the upward trend is statistically significant with *p* = 0.005.

Despite natural variability in the data, which may reflect unmeasured environmental fluctuations or measurement noise, the monotonic rise of the trend line with *D* provides strong confirmation of the theoretical prediction and underscores the value of this parameter in capturing systematic structure in abundance correlations.

## IV. DISCUSSION

## A. Biological Origins of Yield–Depletion Asymmetry

The use of correlations to identify shared dynamical patterns is very popular, despite criticism that has been raised against it in the past [18, 19]. As we have seen throughout this paper, abundance correlations between coexisting species do not reflect niche overlap (NO), but rather the balance between shared responses to environmental fluctuations (which lead to positive correlations) and competition (which lead to negative correlations). In cases where the niche structure involves resource partitioning, as in the model considered here, the key quantity is the difference between the depletion gap ΔΛ and the yield gap ΔΓ.

In our analysis, we assumed that the matrices Γ and Λ are not necessarily equal, and therefore sampled their entries as entirely independent up to the global constraint on the value of NO. By contrast, in the paper of Sireci *et al*. [10] mentioned above, the authors analyzed the same consumer-resource dynamics but assumed that *γ*_*i,j*_ = *λ*_*i,j*_ up to a multiplicative factor. As a result, that study covered only the case *D* = 0, and accordingly, no correlations in consumer abundances were found. Sireci *et al*. [10] thus had to assume that the empirically observed correlations arise from shared responses to environmental factors external to the consumer-resource system, such as temperature, pH, and similar variables. Therefore, our work suggests an alternative interpretation of their results: they can be explained within the consumer–resource framework, provided that genetic distance between microbial strains primarily reflects depletion gap rather than yield gap. This distinction is not only theoretical: empirical yield and depletion profiles often differ substantially, making the YDM framework directly applicable.

Indeed, recent studies have increasingly adopted the assumption that the ratio between yield and depletion is not fixed [5, 13, 20, 21]. This assumption is biologically plausible for several reasons. Even within a single species, the yield-to-depletion ratio can vary substantially with environmental and physiological conditions, such as oxygen availability (aerobic vs. anaerobic metabolism [22]), temperature [23], and metabolic state [24]. Across taxa, physiological strategies also differ: ectotherms like insects and fish often achieve higher conversion efficiency than endotherms like mammals, which invest more energy in maintenance and thermoregulation [25]. Moreover, organisms sometimes deplete resources without direct benefit, for instance, surplus killing by predators [26], destructive foraging by elephants [27], scatter hoarding by rodents [28], or territorial scent-marking [29]. While some correlation between yield and depletion is expected, they are unlikely to be identical. Realistic ecological scenarios likely fall between the limiting case analyzed by Sireci *et al*. [10] and the more general framework considered here. Crucially, once symmetry between yield and depletion is broken, nontrivial correlation patterns emerge.

Furthermore, the empirical data from Crocker *et al*. [5] showed that the yield and depletion profiles (*γ* and *λ*) are not simply related, and this lack of alignment is reflected in the presence of nontrivial abundance correlations, which vanish in the special case *D* = 0.

We focused on the case where the Lotka–Volterra dynamics emerging from the consumer–resource equations involve symmetric interactions, i.e., *α*_*i,j*_ = *α*_*j,i*_. We have also analyzed the two-species case with asymmetric interactions (*α*_*i,j*_ *≠ α*_*j,i*_) and obtained qualitatively similar results. Note that asymmetric Lotka–Volterra dynamics can arise from an underlying consumer–resource system only when Γ *≠* Λ. In diverse communities, such asymmetry may lead to chaotic dynamics with no deterministic steady state, as discussed in [13, 21].

### B. Implications for the role of niche overlap

From a broader theoretical perspective, our findings highlight a conceptual separation between in-niche parameters, those encoded in the reduced competition matrix and governing equilibrium coexistence, and out-of-niche parameters, which control the community’s transient and stochastic dynamics. NO belongs to the former category, whereas the yield–depletion mismatch (YDM) is an example of the latter. This distinction explains why NO alone is insufficient to predict abundance correlations and, more generally, why static coexistence metrics may fail to capture dynamical responses [8].

From an empirical standpoint, these results call for caution in interpreting abundance correlations as direct indicators of NO. While such correlations have been widely used for this purpose, we find that they are strongly influenced by YDM and may bear little relation to NO itself. For practitioners aiming to infer NO from data, our analysis indicates that correlations in instantaneous growth rates, rather than abundances, provide a more reliable proxy, as they more directly reflect shared environmental responses without the delayed confounding effects introduced by competition.

Although our work focuses on abundance correlations, its logic suggests that other out-of-equilibrium properties - invasion probabilities, coexistence robustness, and resilience to environmental disturbances - may likewise depend on out-of-niche parameters rather than on NO. Testing these predictions will require both theoretical extensions and empirical measurements in diverse systems.

Finally, our results have potential implications for understanding the evolutionary process of sympatric speciation. For speciation not to result in the extinction of the ancestral species, the fitness differences between the wild type and the mutant must be small relative to the level of stochasticity in the system, leading to dynamics that are neutral or nearly neutral. As we have shown, there exists a considerable parameter space for such dynamics; that is, many values of Γ and Λ yield *α*_*i,j*_ ≈ 1. We also observed that in these cases, the correlations tend to be negative, implying that the mutant benefits from a storage effect [30, 31] that allows it to invade despite lacking a fitness advantage. This, in turn, may facilitate the accumulation of more and more species. A detailed analysis of such processes can enhance our understanding of the fundamental mechanisms underlying the vast diversity of life observed on Earth.

## Acknowledgments

N.M.S acknowledges support from the Israel Ministry of Science (Italy-Israel cooperation, grant no. 7578) and of the Israel Science Foundation (grant no. 2435/24). We thank Matthieu Barbier, Osnat Gillor and Matteo Sireci for many useful comments and discussions.

## Supplementary A From consumer-resource to Lotka-Volterra

Many mathematical models have been developed to describe deterministic competition between species, with the **generalized Lotka-Volterra (GLV)** model emerging as one of the most widely recognized frameworks. The GLV model has garnered significant praise for its ability to represent complex ecological systems in a canonical and accessible manner. By capturing the general trends in deterministic ecosystem dynamics, the model elucidates emergent behaviors from a minimal set of key parameters [32, 33].

On the other hand, Lotka–Volterra is an “effective model” that defines the level of niche overlap between species but does not address the question of the origin of this overlap. As we have seen throughout this paper, the question of correlations depends not only on the niche overlap itself but also on the mechanistic factors it reflects. Therefore, while we wish to implement the Lotka–Volterra model in which niche overlap is well-defined and transparent, we also seek a more fundamental model from which the Lotka–Volterra equations can be derived. For this purpose, we will use a version of MacArthur’s consumer–resource model.

Another important aspect is the origin of environmental stochasticity in the system. A stochastic element can be added to the Lotka–Volterra equations, but again, its mechanistic origin is unclear. When we use consumer–resource equations, we introduce the stochasticity through the growth rate of the resource, which in turn affects resource availability and, indirectly, the abundance of the consumer. This makes the entire process much more transparent.

### A1. THE GENERALIZED LOTKA-VOLTERRA (GLV) DYNAMICS

The GLV model is governed by the following equation:

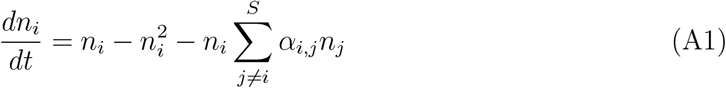

In this formulation:

- *n*_*i*_ represents the population size of species *i*,
- *S* is the number of species,
- The term 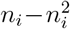 accounts for intrinsic growth and self-limiting effects (logistic growth),
- The term 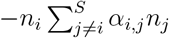 captures inter-species interactions, where *α*_*i,j*_ denotes the interaction matrix.

All in all, the nonlinear interactions are reflected in the ***α*** matrix, whose elements, in the case of three species competition, are

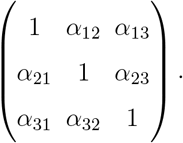

Most of the results presented in this paper concern symmetric ***α*** matrices (*α*_*i,j*_ = *α*_*j,i*_; however, as noted in the main text, we observed the same behavior in asymmetric cases, provided the dynamics admits equilibrium states.

### A2. THE CONSUMER-RESOURCE (CR) MODEL

Our version of MacArthur consumer-resource model is,

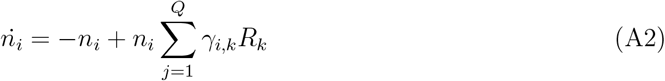

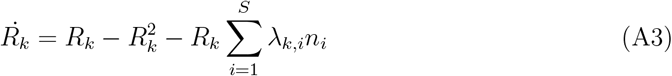

Where:

- *R*_*k*_ is the density of resource *k*,
- *Q* is the number of resources,
- *γ*_*i,k*_ is the yield matrix (how the availability of the resource *k* is translated into the growth rate of the consumer *i*),
- *λ*_*k,ℓ*_ is the depletion matrix (how the presence of the consumer *i* affects the abundance of the resource *k*).

### A3. DERIVATION OF THE GLV MODEL FROM THE CR DYNAMIC

The consumer–resource model is our workhorse in this paper. To derive the corresponding GLV from it, one assumes 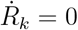, yielding,

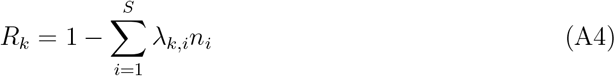

Substituting Eq. (A4) into Eq. (A2), we obtain:

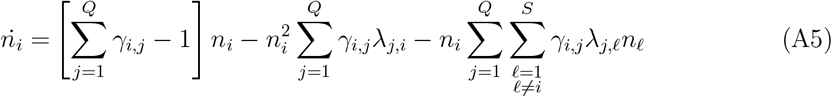

This formulation enables direct comparison between MacArthur’s CR model and the GLV model (Eq. (A1)) by choosing parameters such that:

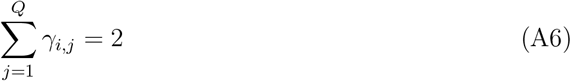

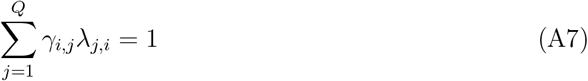

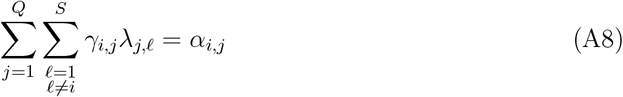

The justification for the assumption 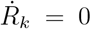 can be given in two ways. One is the assumption that the dynamics of the resource are much faster than those of the consumer; hence, the resource lies on a so-called “fast manifold” and reaches equilibrium before the consumer changes appreciably. However, even if we drop this assumption, as long as an equilibrium state of the system exists, it must satisfy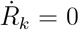, and therefore the consumers will obey the effective Lotka–Volterra equations.

### A4. PARAMETER REDUNDANCY

The constraints shown in Eqs. (A6)-(A8) can be formalized using matrix notation. Let **Γ** be the yield matrix with elements *γ*_*i,j*_, and **Λ** be the depletion matrix with elements *λ*_*j,i*_. Eqs. (A7)-(A8) implies that the interaction matrix ***α*** is given by:

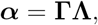

and the linear growth rate of each consumer species is one iff Eqs. (A6) is satisfied, i.e., if each row of **Γ** is constrained to sum to 2.

This mapping, however, is highly redundant: many different pairs of **Γ, Λ** matrices yield the same ***α***. In the deterministic regime, this redundancy is inconsequential, as the resulting GLV dynamics depend only on ***α***. But in the presence of environmental noise, the specific structure of **Γ** and **Λ** influences how fluctuations propagate through the system, making the choice of parameterization meaningful.

#### Example (redundant factorizations)

As an example, let us show how two distinct pairs (**Γ**^(1)^, **Λ**^(1)^) and (**Γ**^(2)^, **Λ**^(2)^) both produce the same GLV interaction matrix

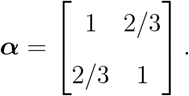

One can easily verify that both **Γ**^(1)^**Λ**^(1)^ = ***α*** and **Γ**^(2)^**Λ**^(2)^ = ***α***, where

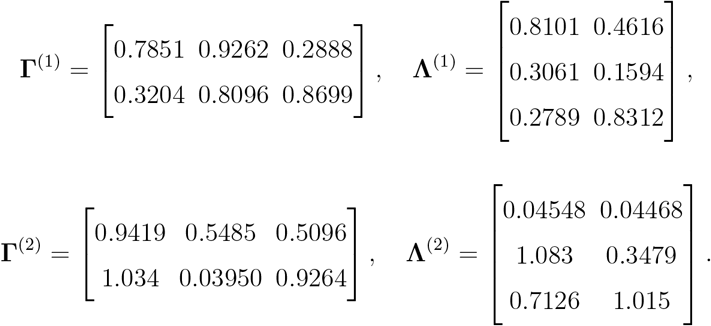

Entries were rounded to four significant figures; each row of **Γ** sums to 2.

### A5. GENERATING YIELD AND DEPLETION MATRICES

Throughout this paper, we fixed the parameter values in the GLV model (thereby also determining the niche overlap) and then searched for many CR systems that, in the limit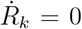, yield the same parameter values, making use of the redundancy. To this end, we employed two methods. Here we provide the details of these two methods.

#### Sampling Method 1 – Exploring Λ Through Null-Space Perturbations

Each row of Γ was generated by drawing a random vector with positive entries and normalizing it so that the row sums to 2:

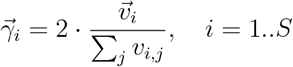

This ensures the constraint (A6).

Given such a Γ, we constructed a compatible Λ by solving:

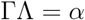

Since Γ is a *S × Q* matrix, the solution for Λ is underdetermined. The Moore-Penrose pseudoinverse Γ^*†*^ provides us with a particular solution:

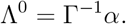

To explore the solution space, we added to this particular solution a random component from the null space of Γ, yielding:

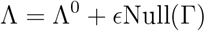

where *ϵ* is a random matrix of appropriate dimensions. This method ensures that all generated Λ matrices satisfy the constraints (A6)-(A8) while allowing for variability due to redundancy in the CR-to-GLV projection.

#### Sampling Method 2 – Constrained Optimization Approach

In this method, we used numerical optimization to generate matrices Γ and Λ that approximately satisfy the condition ΓΛ ≈ *α*.

We jointly optimized the entries of Γ ∈ ℝ^*S×Q*^ and Λ ∈ ℝ^*Q×S*^ by minimizing the following cost function:

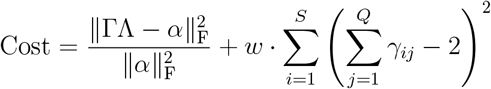

Here, the first term is the normalized Frobenius norm quantifying the reconstruction error between the product ΓΛ and the target interaction matrix *α*. The second term penalizes deviations from the desired row-sum constraint ∑_*j*_ *γ*_*ij*_ = 2, and *w* is a tunable penalty weight.

The optimization was subject to the hard constraint that all elements of Γ and Λ lie within the interval [0, 2]. This was implemented using the Sequential Quadratic Programming (SQP) algorithm within MATLAB’s `fmincon` solver.

## Supplementary B Calculating correlations

To understand how environmental fluctuations propagate through the consumer–resource dynamics and generate correlations in consumer species abundances, we analyze the system near its deterministic equilibrium. Specifically, we linearize the following stochastic differential equations:

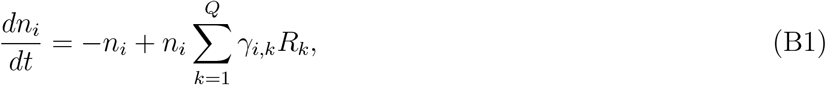

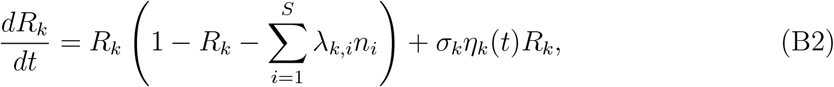

where *n*_*i*_(*t*) and *R*_*k*_(*t*) denote the abundances of consumer *i* and resource *k*, respectively. The matrices *γ*_*i,k*_ and *λ*_*k,i*_ encode the yield and depletion coefficients, and *η*_*k*_(*t*) are independent, zero-mean Gaussian white noise processes, 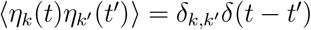.

Our goal is to compute the abundance correlations ⟨*δn*_*i*_*δn*_*j*_⟩ and the growth rate corre-lations ⟨*δr*_*i*_*δr*_*j*_⟩, for consumer species *i* and *j*, at the vicinity of the steady state. In what follows we provide three different methods: a semi-analytic approach, a numerical approach and a full analytic solution for a simple cases.

### B1. THE SEMI-ANALYTIC APPROACH

We decompose each variable into its deterministic equilibrium value and a stochastic fluctuation:

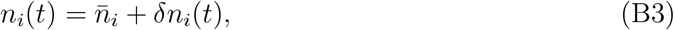

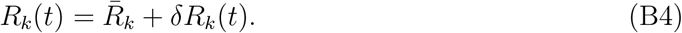

Expanding the dynamics to first order in *δn*_*i*_ and *δR*_*k*_ leads to a linearized system. We collect all fluctuations into a single vector:

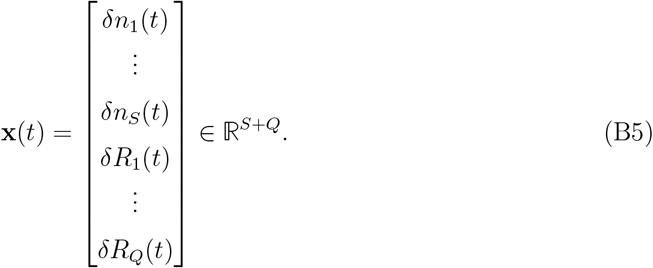

The linearized stochastic dynamics can now be expressed as:

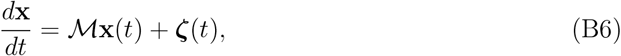

where *ℳ* ∈ ℝ^(*S*+*Q*)*×*(*S*+*Q*)^ is the Jacobian matrix evaluated at equilibrium, and ***ζ***(*t*) encodes the noise terms acting on the system.

#### A. The structure of stochasticity

The noise is assumed to act only on the resource variables and to be proportional to the equilibrium abundance of each resource. Thus, the noise vector takes the form:

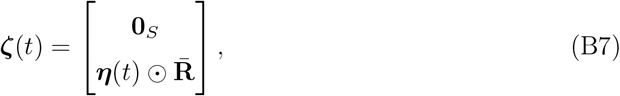

where ***η***(*t*) ∈ ℝ^*Q*^ is a vector of independent white noise processes with

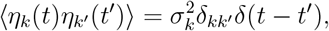

and ⊙ denotes elementwise multiplication.

The noise covariance matrix is then:

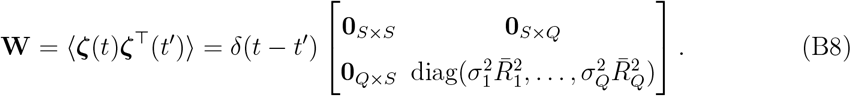

#### B. Abundance Correlations

We denote the steady-state covariance matrix as:

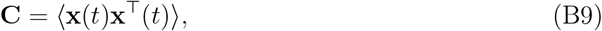

where **x**(*t*) collects both consumer and resource fluctuations. In the steady state, this covariance matrix satisfies the Lyapunov equation:

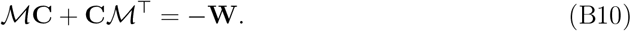

Let *ℳ* be diagonalizable, i.e., *ℳ* = **P**Λ**T** where:

- Θ = diag(*θ*_1_, …, *θ*_*S*+*Q*_) is the diagonal matrix of eigenvalues,
- **P, T** ∈ ℝ^(*S*+*Q*)*×*(*S*+*Q*)^ are the right and left eigenvector matrices satisfying **T**ℳ**P** = Θ and **TP** = **I**.

Then, the covariance between any two components *x*_*m*_ and *x*_*ℓ*_ of the fluctuation vector (consumers or resources) can be written as:

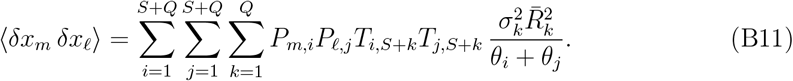

Equation (B11) is a general result: it applies equally to consumer–consumer, resource–resource, and mixed consumer–resource correlations. In particular, the resource–resource covariances ⟨*δR*_*i*_*δR*_*j*_⟩ used later in the growth-rate analysis follow directly from this formula.

#### C. Correlation in Growth Rates

The instantaneous growth rate of species *i* over a small time interval Δ*t* is defined as:

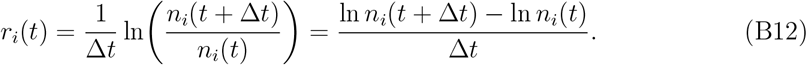

In the limit Δ*t* → 0, this becomes the time derivative of the logarithm of abundance:

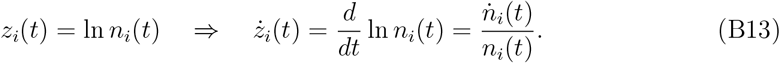

Thus, we define the instantaneous growth rate as:

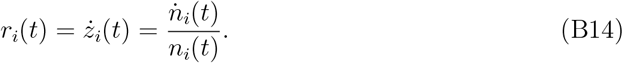

Linearizing the dynamics around the stable equilibrium and assuming that fluctuations in species growth rates are primarily driven by fluctuations in resource levels, we obtain the approximation:

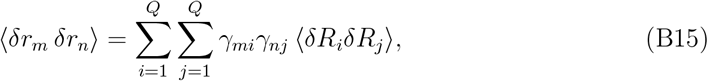

where:

- *γ*_*mi*_ is the entry of the consumption matrix ***γ***, describing the efficiency with which species *m* utilizes resource *i*,
- ⟨*δR*_*i*_*δR*_*j*_⟩ is the covariance of resource fluctuations, obtained from Eq. (B11).

This formulation shows how species that depend on similar resources exhibit correlated growth-rate fluctuations. The magnitude and sign of ⟨*δr*_*m*_*δr*_*n*_⟩ are shaped both by the overlap in resource preferences (via *γ*_*mi*_) and by the resource covariances determined from the general abundance correlation formula.

Calculating the correlations for both abundance and growth rate can also be done numerically by integrating the stochastic differential equations over time. We have compared out numerical with the analytic results and the fits align. This reinforces the robustness of our theoretical framework.

### B2. NUMERICAL INTEGRATION OF SDE

We complement the linear response analysis with direct numerical simulations of the full stochastic system, integrating Eqs. (B1)–(B2) using Stratonovich integration scheme. The continuous-time equations are:

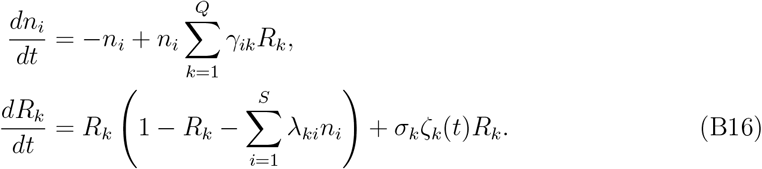

To integrate these equations numerically, we discretize time into steps of size Δ*t*, and implemented,

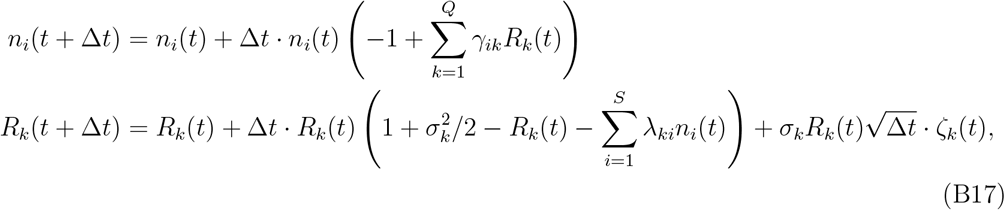

where *ζ*_*k*_(*t*) are independent Gaussian random variables with mean zero and unit variance.

#### A. Parameter Values and Their Role

We used the following parameters in our simulations:

- **Total time** *T* = 10000: the number of full time units simulated.
- **Time resolution** `steps` = 100: number of integration steps per time unit, yielding a time step of Δ*t* = 1`/steps` = 0.01.
- ***σ***_*e*_ = 0.05: strength of environmental noise acting on the resources.
- *N*_*τ*_ = 10: number of resource substeps per consumer step to enforce faster resource dynamics (see below).

##### 1. Time Scale Separation

To mimic biological scenarios in which resource dynamics operate on faster timescales than consumer dynamics, we updated resource abundances more frequently. Specifically, within each consumer time step of size Δ*t*, we performed *N*_*τ*_ = 10 substeps where only the resource variables were updated using Eq. (B17). This introduces an effective time scale separation and better approximates the quasi-steady-state assumption often used in analytical reductions.

##### 2. Measurement of Correlations

After a sufficient transient period, we recorded:

- The time series of species abundances *n*_*i*_(*t*).
- The instantaneous growth rates:
- 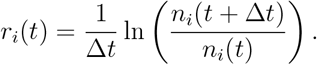

From these, we computed the Pearson correlation coefficients between all species pairs *i, j*:

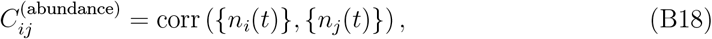

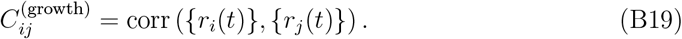

Only the post-transient portion of the simulation was used to compute these statistics.

### B3. NUMERICAL VALIDATION OF THEORETICAL PREDICTIONS

To validate the analytical framework developed above, we performed numerical simulations of the stochastic consumer–resource system with parameter choice *α* = 0.9. The results confirm that the semi-analytic predictions for both abundance and growth-rate correlations are in excellent agreement with direct numerical integration.

#### A. Single Run Example

Figure B1 shows the time series of two consumer species (*n*_1_, *n*_2_) and three resources (*R*_1_, *R*_2_, *R*_3_) over a long simulation. The consumers fluctuate around their steady-state abundances while the resources remain stable, subject to noise-induced fluctuations.

**FIG. B1:**
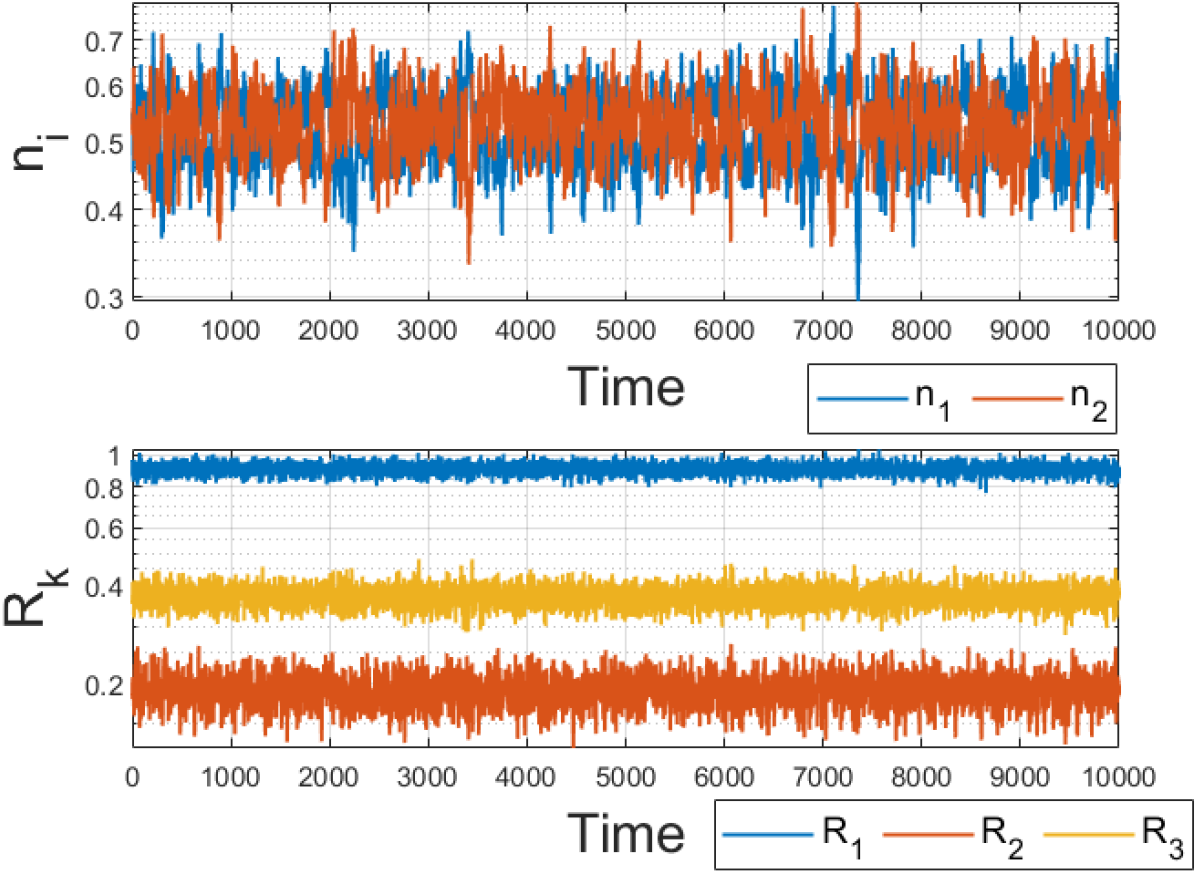
Time series of consumer abundances (*n*_1_, *n*_2_) and resource abundances (*R*_1_, *R*_2_, *R*_3_) from a numerical integration of the stochastic consumer–resource model with *α* = 0.9.

From these trajectories, we computed correlations and compared them with theoretical predictions:

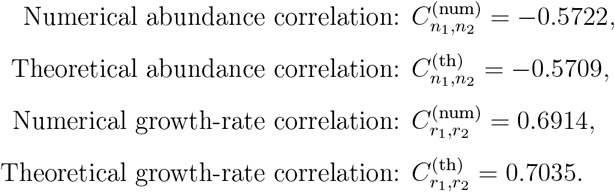

The near-perfect agreement illustrates the robustness of the linear response analysis.

#### B. Ensemble of Trials

To further assess consistency, we repeated the numerical experiment across 100 independent trials with randomized noise realizations. Figure B2 compares the correlations obtained numerically (crosses) with the theoretical predictions (circles). The top panel shows consumer abundance correlations *CR*(*n*_1_, *n*_2_), while the bottom panel shows growth-rate correlations *CR*(*r*_1_, *r*_2_).

**FIG. B2:**
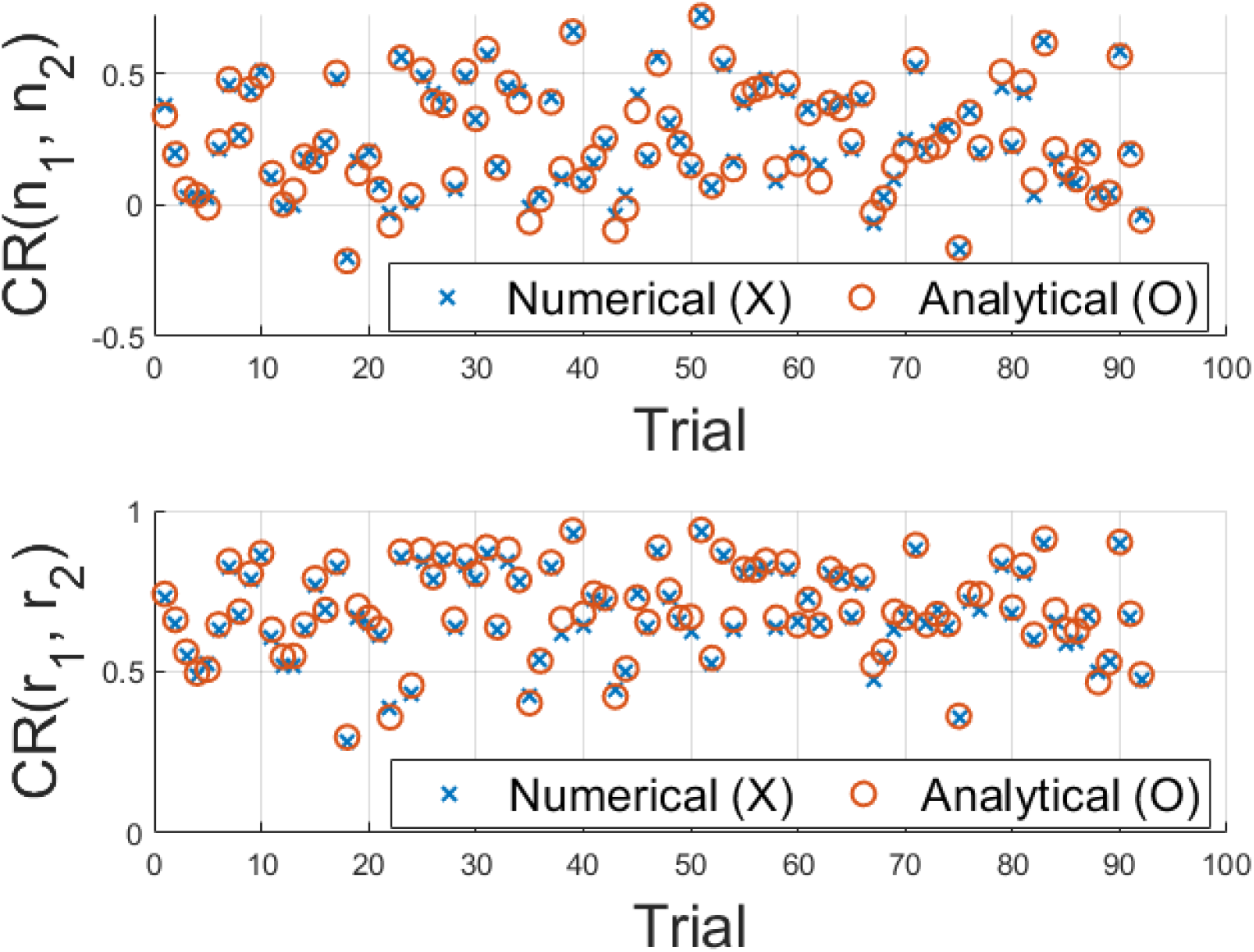
Comparison of numerical (X) and analytical (O) correlations across 100 independent trials. Top: abundance correlations *CR*(*n*_1_, *n*_2_). Bottom: growth-rate correlations *CR*(*r*_1_, *r*_2_). The close alignment across trials demonstrates the accuracy of the theoretical predictions.

## Supplementary C The Yield-Depletion Mismatch (YDM) metric

As shown in the main text, the correlation function is determined by the matrices Γ and Λ. Our aim, therefore, is to identify a clear and tractable metric that captures the key features relevant to the correlation function.

To guide a theory-based conjecture, we first consider a simple illustrative case that admits a transparent analytic solution. Specifically, we select Γ and Λ matrices that generate the effective interaction matrix,

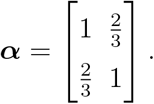

We require Γ and Λ such that ***α*** = ΓΛ and each row of Γ sums to 2. To allow for a fully analytic solution, we focus on a system of two species and two resources, with the additional condition that the diagonal elements of Λ are equal. Concretely, the two matrices take the form

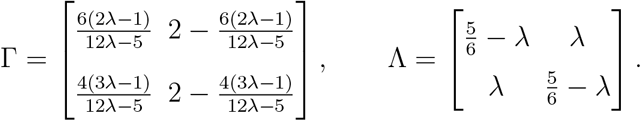

We then calculated the abundance correlation using the method described in Eq. (B11) of Supplementary B. Because the matrices take such a straightforward form, the abundance correlations between the consumer species can be expressed in a concise analytic formula,

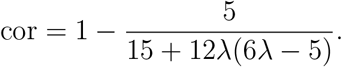

Next, we calculated the YDM parameter *D* as described in the main text. For every pair of species *i* and *j*,

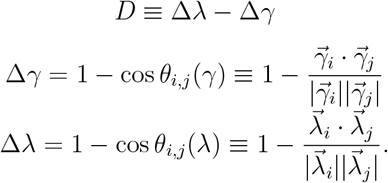

Here, the vector 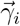 is the *i*-th row of the yield matrix Γ, and 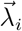 is the *i*-th column of the depletion matrix Λ; |*·* | denotes the Euclidean norm.

For the matrices in our two-species, two-resource system, the explicit expression for *D* is

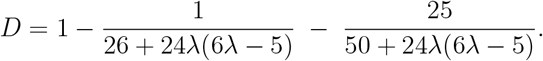

Clearly, cor and *D* are not the same quantity. Still, a glance at Fig. C1 shows that these two functions share the same qualitative shape, attain the same peak value, and cross zero at identical values of *λ*. In other words, the YDM parameter provides a faithful estimator for the correlation derived directly from the stochastic dynamics. Importantly, this agreement is not limited to the specific example above: for any choice of *α*, the peak and zero-crossings of cor and *D* coincide.

As noted in the main text, another advantage of *D* is that it is a bounded, normalized metric: since 0 ≤ Δ_*γ*_, Δ_*λ*_ ≤ 2, it follows that −2 ≤ *D* ≤ 2. Consequently, *D* is scale-free and enables consistent comparisons across systems.

**FIG. C1:**
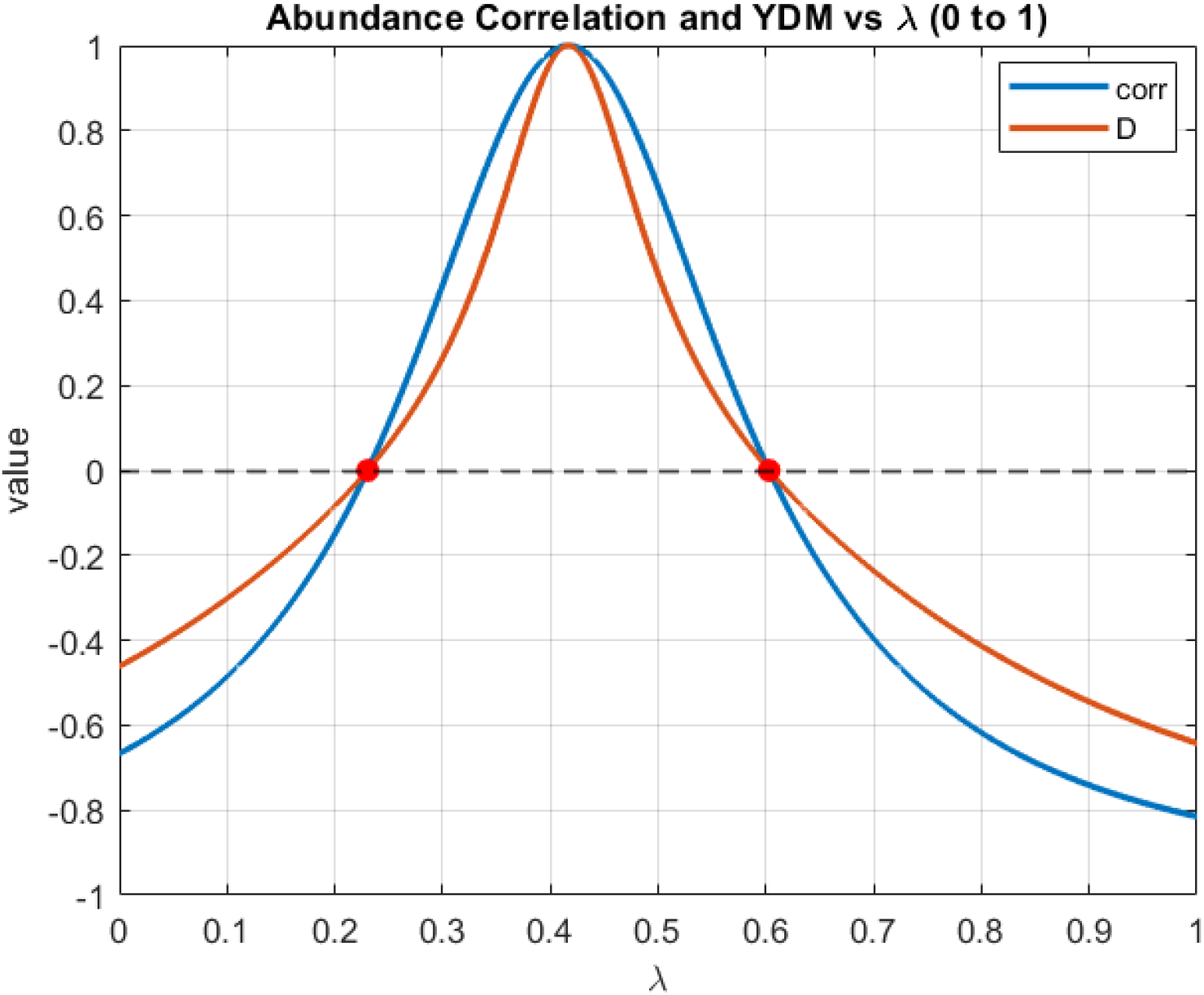
Comparison of the abundance correlation cor(*λ*) and the YDM parameter *D*(*λ*). Both functions exhibit the same qualitative shape, share the same peak value, and cross zero at the same points.

## Supplementary D Empirical Results

### D1. DETAILS OF CROCKER’S EXPERIMENT

Crocker et al. [5] sought to reduce the complexity of microbial community dynamics by identifying *functional guilds*, groups of strains with similar metabolic traits that respond cohesively to environmental changes. Their central idea was that although microbial communities contain many species, the number of distinct “functional roles” may be far smaller. If these roles can be experimentally identified, community dynamics can be coarse-grained from species level to guild level.

To achieve this, they constructed a synthetic community of 20 soil bacterial strains and characterized each strain in monoculture across 10 carbon sources (arabinose, butyrate, deoxyribose, glucuronic acid, glycerol, mannitol, mannose, melibiose, propionate, and raffinose). From these monocultures they measured:

- the exponential growth rate on each carbon source (via OD measurements),
- the biomass yield per unit carbon,
- and the resource uptake rate, inferred as *r*_*i,α*_ = *g*_*i,α*_*/η*_*i,α*_ from growth and yield.

These measurements provided a growth matrix *G* describing the metabolic profile of each strain. By clustering rows of *G*, Crocker et al. defined functional guilds, i.e. sets of strains with overlapping metabolic niches.

The full 20-strain community was then assembled and inoculated into 32 distinct environments, each defined by a random subset of the 10 carbon sources (with total carbon fixed at 25 mM). Each environment was propagated through 9 serial batch cycles of 48 hours growth followed by a 1:10 dilution into fresh medium. At the end of each cycle, absolute strain abundances were determined using 16S sequencing with a spike-in calibration.

From these experiments Crocker et al. demonstrated that guild cohesion depends on the timescale of environmental fluctuations: on short timescales, strains within a guild fluctuate positively together (shared responses to nutrient pulses), whereas on longer timescales, competitive interactions dominate and abundances of guild members become negatively correlated.

#### A. D2. Use of Crocker’s Dataset in Our Work

While Crocker et al. focused on guild-level responses to environmental fluctuation timescales, our analysis addressed a different question: whether pairwise abundance correlations reflect niche overlap (NO) or instead the *yield–depletion mismatch* (YDM).

For this purpose, we drew directly on the dataset deposited in their public repository (https://doi.org/10.17605/OSF.IO/J8S2V). In particular, we used three processed tables:

- `Gamma.xlsx(Find actual file name)` — growth (yield) matrix from monocultures,
- `Yield.xlsx(Find actual file name)` — biomass yields on each carbon source,
- `strain_abundance_matrix.csv(Find actual file name)` — time series of strain abundances across 9 cycles in 32 environments.

Using the monoculture data, we computed functional dissimilarities for each pair of strains: the yield distance Δ*γ* (cosine distance between rows of Γ) and the depletion distance

Δ*λ* (cosine distance between columns of Λ). Using the community time series, we calculated abundance correlations (Spearman’s *ρ*) across replicates and environments.

This allowed us to test our theoretical prediction that correlations are governed not by NO itself, but by the asymmetry between depletion and yield profiles, quantified by our metric

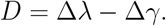

Thus, while Crocker’s analysis emphasized coarse-graining dynamics into guilds, our reanalysis repurposed the same experimental system to uncover the mechanistic role of yield– depletion mismatch in driving co-fluctuations.

### D2. ROBUSTNESS ANALYSIS

To test the robustness of our empirical findings, we systematically explored how the relationship between the yield–depletion mismatch parameter *D* and abundance correlations depends on three filtering parameters:

- `top_n_sp`: Number of top-abundant species per environment included in the analysis (5, 10, or 15)
- `rep_start`: The first batch replicate to include (e.g., starting from replicate 1, 2, or 3 to exclude early-time transients)
- `min_count`: Minimum number of environments in which a pair must co-occur to be included in the average (2, 5, or 10)

We applied these filters within the analysis script, which performs the following steps:

1. Loads species abundance trajectories and monoculture yield/depletion data.
2. For each environment, computes relative abundance time series for the top *N* most abundant species.
3. Calculates Spearman correlations between all pairs of selected species.
4. Computes the corresponding yield–depletion mismatch *D* = Δ*λ* − Δ*γ* for each pair.
5. Bins data by unique *D* values (only including those with ≥`min_count` replicates) and computes mean correlation and standard error of the mean (SEM) per bin.
6. Performs a weighted linear regression using 1/SEM as weights to assess the significance of the trend.

### D3. SUMMARY OF RESULTS

The table below summarizes the regression results for each combination of filtering parameters. For `top_n_sp` = 10 and 15, we observe a consistent, statistically significant positive slope in the relationship between *D* and abundance correlation, with *p*-values well below 0.05. In contrast, results for `top_n_sp` = 5 show weaker and nonsignificant trends, due to insufficient sample size and increased noise.

**TABLE D1:** Weighted linear regression of abundance correlation vs. *D*, under various filtering conditions.

| Top N | Min Count | Replicates | Fit Equation | p-value |
| --- | --- | --- | --- | --- |
| 10 | 2 | 1-9 | $2.5833 \cdot D - 0.0051$ | 0.00327 |
| 10 | 2 | 2-9 | $2.6915 \cdot D - 0.0290$ | 0.00132 |
| 10 | 2 | 3-9 | $2.5729 \cdot D + 0.0069$ | 0.00480 |
| 10 | 5 | 1-9 | $2.5005 \cdot D - 0.0569$ | 0.00722 |
| 10 | 5 | 2-9 | $2.5624 \cdot D - 0.0656$ | 0.00581 |
| 10 | 5 | 3-9 | $2.2118 \cdot D - 0.0522$ | 0.01410 |
| 10 | 10 | 1-9 | $2.5981 \cdot D - 0.0790$ | 0.01550 |
| 10 | 10 | 2-9 | $2.4302 \cdot D - 0.0820$ | 0.02890 |
| 10 | 10 | 3-9 | $2.1173 \cdot D - 0.0781$ | 0.05350 |
| 15 | 2 | 1-9 | $1.3832 \cdot D + 0.0217$ | 0.00643 |
| 15 | 2 | 2-9 | $1.4242 \cdot D + 0.0164$ | 0.00605 |
| 15 | 2 | 3-9 | $1.4101 \cdot D + 0.0117$ | 0.00622 |
| 15 | 5 | 1-9 | $1.3931 \cdot D + 0.0211$ | 0.00899 |
| 15 | 5 | 2-9 | $1.4449 \cdot D + 0.0141$ | 0.00548 |
| 15 | 5 | 3-9 | $1.3473 \cdot D + 0.0153$ | 0.01120 |
| 15 | 10 | 1-9 | $1.3961 \cdot D + 0.0208$ | 0.00921 |
| 15 | 10 | 2-9 | $1.4475 \cdot D + 0.0139$ | 0.00566 |
| 15 | 10 | 3-9 | $1.3445 \cdot D + 0.0156$ | 0.01180 |
| 5 | 2 | 1-9 | $2.4353 \cdot D - 0.2048$ | 0.32500 |
| 5 | 2 | 2-9 | $2.8804 \cdot D - 0.2195$ | 0.25400 |
| 5 | 2 | 3-9 | $3.1423 \cdot D - 0.2053$ | 0.23800 |
| 5 | 5 | 1-9 | $2.7975 \cdot D - 0.2013$ | 0.35000 |
| 5 | 5 | 2-9 | $3.3280 \cdot D - 0.2274$ | 0.24600 |
| 5 | 5 | 3-9 | $4.9231 \cdot D - 0.2606$ | 0.13900 |
| 5 | 10 | 1-9 | $3.1753 \cdot D - 0.3252$ | 0.38100 |
| 5 | 10 | 2-9 | $4.1148 \cdot D - 0.3683$ | 0.16900 |
| 5 | 10 | 3-9 | $4.2064 \cdot D - 0.3786$ | 0.14400 |

These findings confirm that the predictive power of *D* is robust across filtering schemes and supports our theoretical prediction that abundance correlations are governed by yield–depletion asymmetry rather than total functional dissimilarity.

## Notes

### Competing Interest Statement

The authors have declared no competing interest.

